# NWB:N 2.0: An Accessible Data Standard for Neurophysiology

**DOI:** 10.1101/523035

**Authors:** Oliver Rübel, Andrew Tritt, Benjamin Dichter, Thomas Braun, Nicholas Cain, Nathan Clack, Thomas J. Davidson, Max Dougherty, Jean-Christophe Fillion-Robin, Nile Graddis, Michael Grauer, Justin T. Kiggins, Lawrence Niu, Doruk Ozturk, William Schroeder, Ivan Soltesz, Friedrich T. Sommer, Karel Svoboda, Ng Lydia, Loren M. Frank, Kristofer Bouchard

## Abstract

Neurodata Without Borders: Neurophysiology (NWB:N) is a data standard for neurophysiology, providing neuroscientists with a common standard to share, archive, use, and build common analysis tools for neurophysiology data. With NWB:N version 2.0 (NWB:N 2.0) we made significant advances towards creating a usable standard, software ecosystem, and vibrant community for standardizing neurophysiology data. In this manuscript we focus in particular on the NWB:N data standard schema and present advances towards creating an accessible data standard for neurophysiology.

## 1 Introduction

### Motivation

Brain function is produced by the coordinated activity of multiple neuronal types that are widely distributed across many brain regions. Neuronal signals are acquired using extra- and intracellular recordings, and increasingly optical imaging, during sensory, motor, and cognitive tasks. Neurophysiology research generates large, complex and heterogeneous datasets at terabyte scale. The data size and complexity is expected to continue to grow with the increasing sophistication of experimental apparatuses. Lack of standards for neurophysiology data and related metadata is the single greatest impediment to fully extracting return-on-investment from neurophysiology experiments, impeding interchange and reuse of data and reproduction of derived conclusions. This gap motivated the launch of the Neurodata Without Borders: Neurophysiology (NWB:N) data standards project. The goal of NWB:N is to develop a standardized format and methods for neurophysiology data and metadata.

### Background

The first NWB:N 1.0.x standard was the result of a 1 year pilot project in 2015^12^. As part of this pilot, neurophysiologists and software developers met during two hackathons to create a common data format for recordings and metadata of cellular electro- and optical physiology experiments (Fig. 1, top). Despite the important advances that NWB:N 1.0 made towards creating a neurophysiology data standard, the standard was not easily accessible to users. To enhance broad adoption, a sustainable software and community strategy and easy-to-use, high-level application programming interfaces (APIs) were desperately needed. Here we describe NWB:N 2.0, a modern ecosystem for data standardization and accessible data standard for neurophysiology.

### A Brief History of NWB:N 2.0

The development of the second version of NWB:N began in Janurary 2017 with the start of the Kavli funded NWB4HPC project. The goal was to develop infrastructure and algorithms to enable data-driven discovery and dissemination on high-performance computing systems for the BRAIN Initiative (Fig. 1, bottom). One main goal of the project was to develop the next version of NWB:N to enhance its adoption, with an initial focus on high-level APIs for read, write, and extension of the original NWB:N 1.0.x standard. This standard represented a critical first step toward a unified framework for neural data, but it became clear that in order to achieve these goals we needed an advanced software architecture, a well-articulated data standards ecosystem, an open community software strategy, and advancements to the NWB:N data standard itself. Under leadership of K. Bouchard, O. Ruebel, and A. Tritt (LBNL), in collaboration with F. Sommer, J. Teeters et al. (UCB), the LBNL team developed a modern software strategy for NWB:N, identified and implemented critical changes to the NWB:N data standard, and identified, created, and separated the core components of the NWB:N ecosystem, i.e., the specification language, standard schema, data storage, and data APIs (Sec. 2). Throughout both this early phase of the project and the subsequent development of NWB:N 2.0, the Frank and Chang laboratories at the University of California at San Francisco (UCSF) provided additional funding support to the LBNL team as well as feedback and use-cases.

**Figure 1.**
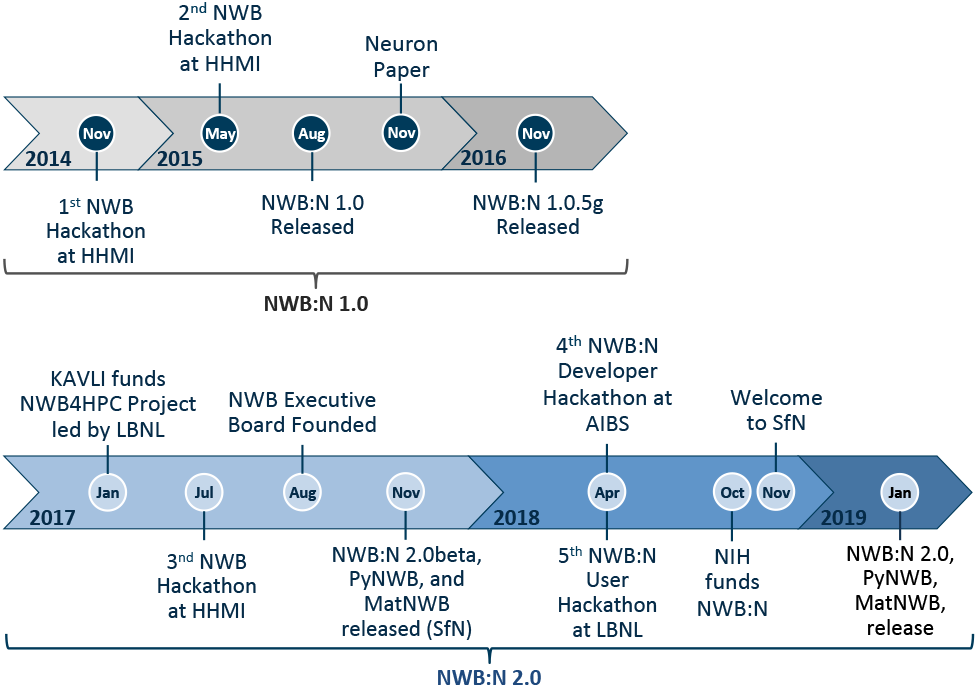
Overview of the history of the NWB:N project.

In July 2017 the 3rd NWB:N hackathon was held at HHMI Janelia, marking the beginning of the second main phase of the NWB:N 2.0 endeavour. The goal of the hackathon was to review the team’s proposal and progress towards NWB:N 2.0 and to create a governance structure for NWB:N. As a result of the hackathon, the NWB:N Executive Board was established in August 2018 and NWB:N 2.0 was officially established as a community project, including the organization and public release of all sources via the NeurodataWithoutBorders GitHub organization and creation of open community channels via Slack and GoogleGroups. Driven by these advances, the user and developer community around NWB:N began to grow as well as engagement with industry partners. Kitware Inc. contributed significantly to the design of continuous integration processes for NWB:N^4^. Vidrio Technologies, in collaboration with the LBNL team, led development of the MatNWB Matlab API for NWB:N. The Allen Institute for Brain Science (AIBS) as a major early adopter of NWB:N also contributed significantly to the development of NWB:N 2.0. A first beta version of NWB:N 2.0, including first full versions of the PyNWB (Python) and MatNWB (Matlab) APIs, was then released in November 2017 in conjunction with the annual Society for Neuroscience (SfN) conference.

With the NWB:N 2.0 Beta release, the third main phase of the NWB:N 2.0 project began, focused on refinement and continued development of NWB:N 2.0 based on feedback and collaboration with a broad range of early adopters and beta testers and the broader community. To facilitate development and integration across the developer community, the 4th NWB:N developer hackathon was held at AIBS on April 3-6, 2018^1^. To enhance user engagement and adoption, a 5th NWB:N user training hackathon was held at LBNL on April 26-27, 2018^5,6^. Based on the feedback from these hackathons and continued feedback and experience from early adopters, development of NWB:N 2.0 then continued through the remainder of 2018. Throughout this process, the broader NWB:N community contributed code and feedback via GitHub, Slack, GoogleGroup, and email.

In October 2018, NIH announced its support to continue development of the NWB:N data standard as part of a BRAIN Initiative grant led by Dr. Ruebel (LBNL) and Dr. Ng (AIBS). In November 2018 the NWB:N 2.0 schema was then finalized and development-focus switched to bug fixes and stabilization of the APIs and tools. The full release of NWB:N 2.0, including the new data standard schema, PyNWB and MatNWB APIs, tools (e.g., nwb-docutils), and supporting documents, was then announced in January 2019.

### Contributions

In this manuscript we present the advances in the NWB:N 2.0 data standard schema compared to the previous NWB:N 1.0.6 standard. In Sec. 2 we first provide a high-level overview of the NWB:N 2.0 ecosystem and highlight specific changes as they relate to the schema. In Sec. 3 we discuss technical advances in the NWB:N 2.0 data standard with a focus on new general capabilities, e.g., support for data referencing, tables, ragged arrays or compound data types. We then discuss in Sec. 4 the application of these new methods in practice to improve storage and management of neurophysiology data as part of the NWB:N 2.0 data standard. Finally, in Sec. 5 we conclude with a discussion of the current NWB:N community and future directions.

## 2 Overview of the NWB:N 2.0 Ecosystem

NWB:N 2.0 is more than just a file format; it defines an ecosystem for standardization of neurophysiology data. In the following we provide a brief overview of the different core components of the NBW:N ecosystem and highlight the main changes compared to NWB:N 1.0.x. In this manuscript we focus on advances in the data standard schema (Sec. 3 and 4). Advances in the software strategy, APIs, schema language and storage will be the focus of other publications.

### 2.1 Specification Language

To support the formal and verifiable specification of neurodata file formats, NWB:N defines and uses the NWB:N specification language^7^. The specification language defines formal structures for describing the organization of complex data using basic concepts, e.g., groups (similar to folders), datasets (n-D arrays), attributes (metadata objects on groups and datasets), and links (to groups/datasets/files). The specification language also enables users to define extensions to the NWB:N format to support the integration and storage of currently unsupported data with NWB:N. NWB:N 2.0 includes a substantial overhaul of the original 1.0.x specification language and provides advanced APIs for creating, reading, and writing specification language documents and tools to generate Sphinx documentation from specifications. See the schema language^7^ and PyNWB^10^ documentations for details.

### 2.2 Data Standard Schema

Using the NWB:N specification language, the NWB:N standard schema formally specifies the organization of neuroscience data^8^. The standard schema provides a verifiable, computer and human readable document that governs the NWB:N format. As a result, the format specification is central to the development of APIs and codes compliant with the NWB:N data standard.

The NWB:N data standard uses a modular design in which all main semantic components of the format have a unique `neurodata_type` (similar to a class in object-oriented design). This allows for reuse and extension of types through inclusion and inheritance. All datasets and groups in the format can be uniquely identified by either their name and/or type. At a high level, data are organized in an NWB:N file in the following main groups:

- `/acquisition` for storage of data streams recorded from the system, e.g., recordings from electro- and opto-physiology or behavioral tracking systems.
- `/processing` for storage of standardized processing modules, often as part of intermediate analyses required before scientific analysis, e.g., results from spike sorting, signal filtering, or image processing.
- `/intervals` for storage of experimental intervals, e.g., experimental epochs or trials.
- `/stimulus` for storage of stimulus data.
- `/general` for storage of experimental metadata, e.g., protocol, notes or device descriptions.
- `/analysis` for storage of lab-specific and custom scientific analysis data.

Neural data typically involves measurements taken over time. Thus, the NWB:N data standard is designed around the concept of `TimeSeries`, a generic neurodata_type for storing time series data that is extended via sub-classing to account for different storage requirements and data modalities (e.g., `ElectricalSeries` for electrophysiology or ImageSeries for optical imaging). The full sources of the NWB:N 2.0 schema are available as YAML online using an open BSD license model. For further details see Sec. 3 and 4 as well as the NWB:N data standard documentation^8^.

### 2.3 Data Storage

The role of the data storage backend is to map NWB:N primitives (Groups, Datasets, Attributes, Links etc.) to storage. Currently NWB:N uses the Hierarchical Data Format (HDF5) as its primary file-based storage mechanism^13^. HDF5 is a mature data format standard that is widely supported across programming languages (e.g., C, C++, Python, MATLAB, R among others) and tools (e.g. HDFView, VisIt, ParaView, Jupyter and many others). Within a single file, HDF5 supports complex data organizations analogous to a file system in which Groups and Datasets correspond to directories and files. This powerful approach enables organization of large-scale and complex data and metadata within a single file. The data modeling primitives of the NWB:N specification language largely mirror the primitive types in HDF5 so that the mapping between the schema and HDF5 storage is largely 1-to-1^9^. Overall, HDF5 is an excellent choice for storage, sharing and transfer of large scientific data. However, the NWB:N community recognizes that as the applications of the NWB:N data standard grow, requirements and needs for data storage may change and depending on the specific use and application area, other storage backends may be preferable, e.g., databases, RDD, JSON, or others. As such, one main consideration in the design of the NWB:N 2.0 software strategy was to insulate the data storage as much as possible from the schema and to provide APIs that will facilitate the integration of new storage methods with the NWB:N ecosystem in the future.

### 2.4 Data API

The role of data API(s) is to facilitate easy and efficient interaction with neuroscience data stored in the NWB:N data format, e.g., for reading, writing, querying, and analyzing neurophysiology data. A data API should provide a stable and usable interface for programmatic use and development of new applications while insulating developers and users from implementation details related to the specification language, format specification, and data storage.

Development of advanced data APIs for NWB:N files and the specification language has been a central focus of the development of NWB:N 2.0. The newly developed PyNWB^10^ (Python) and MatNWB^3^ (Matlab) APIs both provide easy-to-use representations of NWB:N 2.0 neurodata types for programmatic use and enable the mapping of these representations to/from data storage based on the NWB:N format specification. Both APIs also support read/write of custom format extensions. PyNWB further provides functionality for creating schema extensions and defines abstractions to enable the creation of custom data storage backends for NWB:N. We refer the interested reader to the API documentation for detail^3,10^.

## 3 Methods

In the following we describe core technical advances in the NWB:N 2.0 data standard. We will then describe the applications of these methods in practice to enhance management and storage of neurophysiology data in Sec. 4.

### 3.1 Base Data Types

In order to define common functionality and ease future evolution and extension of functionality, NWB:N 2.0 defines core base types for typed groups, datasets, and primary data groups. Beyond specification of common content as part of the schema, a main role of the base types is to allow APIs to implement shared functionality as part of the base types.

`NWBContainer` defines the common base type for all groups with an assigned neurodata_type in the NWB:N 2.0 data standard schema. The specification of `NWBContainer` is minimal and mainly requires a help attribute.

`NWBData` serves a similar function to `NWBContainer` but instead of for groups it defines the common base type for all datasets with an assigned neurodata_type in the schema.

`NWBDataInterface` extends `NWBContainer` and serves as base type for primary data (e.g., experimental or analysis data). In the schema it is used to distinguish between non-metadata data containers and metadata containers. NWBDataInterface serves as base type for all primary data types in NWB:N 2.0, including the `TimeSeries` base type.

### 3.2 Supporting Explicit Data Referencing

The ability to cross-reference data is essential for neurophysiology; multiple sources of data are typically acquired together and analyzed in context of each other. For example, measurements from multiple devices and brain regions as well as external stimuli and behavior.

#### Links

To enable cross-referencing of diverse data, NWB:N 1.0. x supported the creation of links, and the ability to specify and create links has been further enhanced in NWB:N 2.0. Similar to soft links on a file system, links allow users to include existing datasets or groups inside a group by linking to the existing object.

#### Object/Region References

Links are mainly useful in cases where we need to reference single objects, however, their utility is limited in cases where we need to store large collections of references. To address this challenge, NWB:N 2.0 adds support for object references. Object references behave like links but instead of being stored as independent objects as part of the content of a group, object references are stored as values of datasets or attributes (i.e., they are part of the data type). This approach supports management of large collections of references as part of multi-dimensional arrays. NWB:N 2.0 also adds support for region references, which are similar to object references but support linking to subsets of datasets. The ability to store object- and region references has been central to enable many of the advancements in the NWB:N 2.0 data standard schema.

### 3.3 Supporting Compound Data Types

Compound data types are essentially a struct, i.e., the data type is a composition of several primitive types. In practice, this is useful to support complex datatypes, e.g., for storage of complex numbers, vectors or tensors as elements of datasets as well as to create table-like data structures (described next in Sec. 3.4). With regard to the NWB:N specification language, compound types are defined via a list of sub-types each with a name, dtype, and description (Fig. 2).

**Figure 2.**
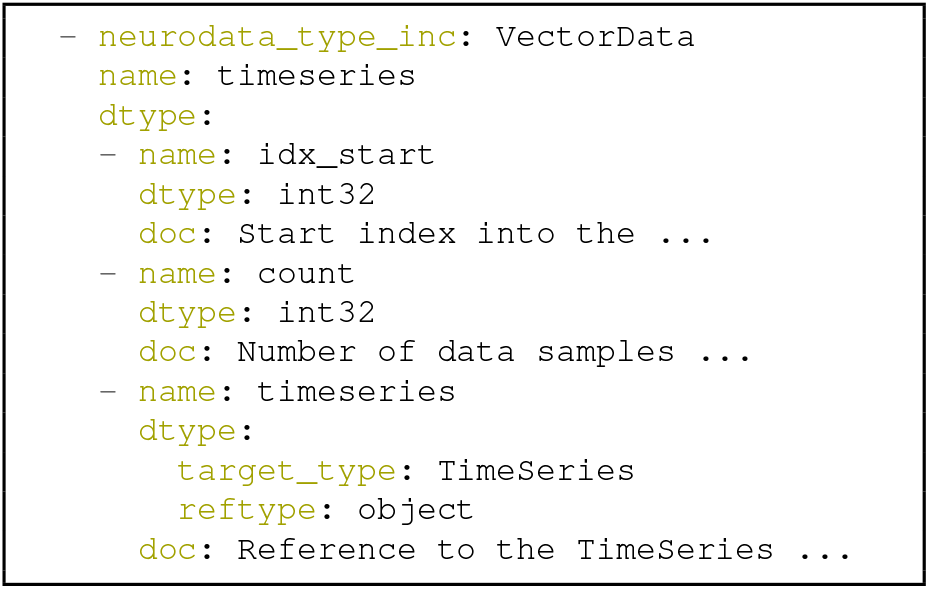
Example illustrating the definition of a compound data type used as part of the management of time intervals (described later in Sec. 4.1.5) to store the mapping of explicit time interval to specific time series. The compound data type stores the integer start index and count along with an object reference to the `TimeSeries` object.

### 3.4 Supporting Data Tables

Data tables are a natural way to organize large collections of related data/metadata. In the context of databases, we typically distinguish between row-based and column-based table storage. To support a wide variety of use cases, NWB:N 2.0 adds support for row-based, column-based, and hybrid approaches for storing tables. In practice, the optimal approach to storing tables depends on the structure, size, and most common read/write access patterns of a table. Broadly speaking, row-based tables generally work well for tables with potentially large numbers of rows, small number of fixed, *a priori* known columns, and primarily row-oriented read/write access. In contrast, column-based tables generally work best for tables with arbitrary (and potentially dynamic) number of columns and primarily column-oriented read/write access.

#### Row-based Tables

Using the concept of compound data types, we can define a data table via a one-dimensional dataset with a compound data type. Here, each element in the data array represents one row and the components of the compound data type describe the table columns (Fig. 3a).

The advantage of row-based tables is that they provide easy access to rows and can be expressed via a single dataset. This allows row-based tables to support: i) referencing of sets of rows via a single region-reference, ii) addition of rows by appending to a single dataset, and iii) fast read of individual rows of a table. Conversely, a main disadvantage of row-based tables is lack of efficient support for column-based operations, e.g.: i) extracting the values of a single column requires reading the full table, ii) referencing of columns is not supported natively, and iii) appending columns requires changing the compound data type and is, hence, not possible dynamically and requires the creation of extensions to the data standard.

**Figure 3.**
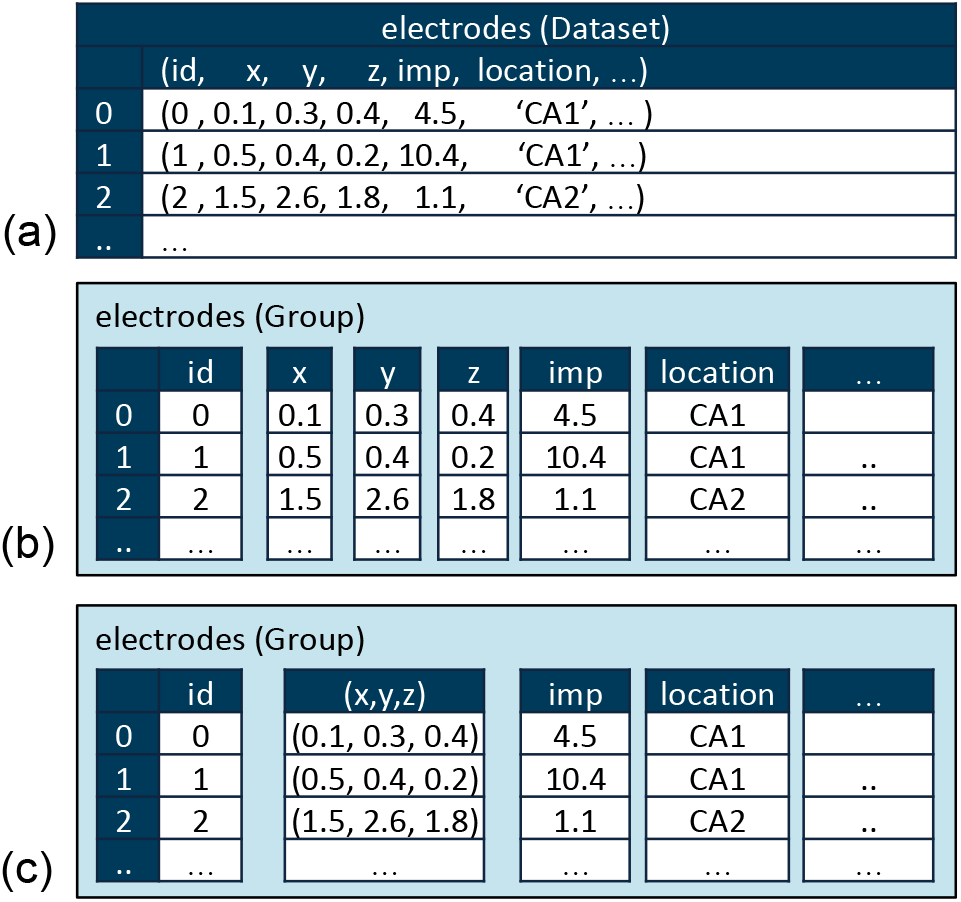
Example of a (a) row-based, (b) column-based and (c) hybrid table storage scheme. See Appendix A for example specifications of these three tables.

#### Column-based Tables

As illustrated in Fig. 3b, NWB:N 2.0 supports column-based tables via the new, reusable type Dynamic Table that stores for each table column a separate VectorData dataset, while the first dimension of each of the column datasets must have the same length as the total number of rows in the table.

The advantage of column-based tables is that they provide easy access to columns and can be more easily extended. This allows column-based tables to support: i) referencing of single columns via simple links or object references, ii) dynamic addition of new columns without the need for extensions to the data standard and iii) easy and efficient read/write of individual columns, and iv) potential to customize data-layout and compression on a per-column basis. Conversely, a main disadvantage of column-based tables lies in the fact the number of data objects required grows linearly with the number of table columns and the resulting overhead for row-based operations, e.g. read/write/append of a single row requires access to multiple datasets. As column-based tables do not natively support row-based references, NWB:N 2.0 provides the dataset type `DynamicTableRegion`, which stores the integer indices of the relevant rows as well as an object reference to the corresponding `DynamicTable`.

#### Hybrid Tables

In the context of `DynamicTable`, NWB:N 2.0 also supports the use of compound type datasets as part of individual table columns. This approach allows for great flexibility to create hybrid row/column table stores. In practice, this approach can be useful, for example, to optimize row-based access to select columns in a dynamic table that are typically accessed in conjunction, e.g., the (*x,y,z*) position of a neuron (Fig. 3c).

**Figure 4.**
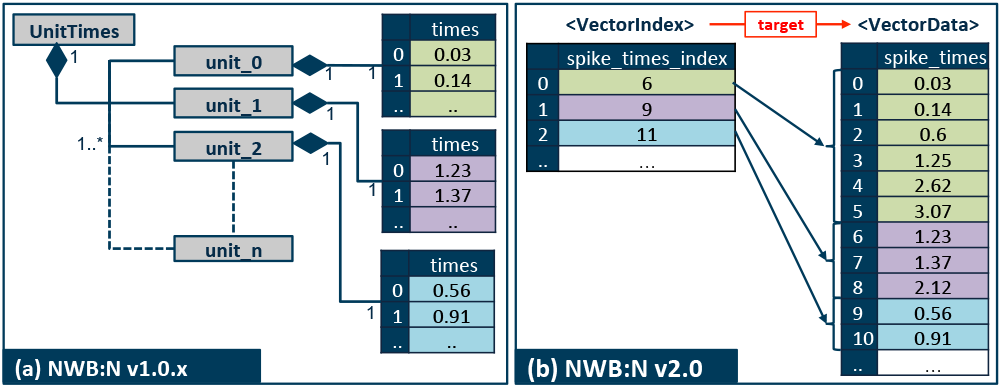
Illustration showing the use of (a) implicit ragged arrays as used in NWB:N 1.0.x, and (b) region-based ragged arrays used in NWB:N 2.0 to store the times of spikes from single units (putative single neurons).

### 3.5 Supporting Ragged Arrays

Ragged arrays (a.k.a., jagged arrays) are arrays where each element of the array is itself an array of variable length. In neurophysiology the need for ragged arrays arises commonly during feature extracting (e.g., detection of spikes or ROIs), where each feature is described by a variable-length vector, e.g., a pixel mask or spike times. Unfortunately, many common data storage solutions do not natively support ragged arrays. Using the NWB:N primitives we can, however, model ragged arrays via implicit ragged arrays (NWB:N 1.0.x) and region-based ragged arrays (NWB:N 2.0) (as well as dense arrays, as discussed in Appendix B). Here we focus on simply-ragged arrays where the array is ragged along the second dimension only and does not itself contain ragged arrays.

#### Implicit Ragged Arrays

The original NWB:N 1.0.x schema used combinations of groups and datasets to store large collections of variable-size arrays. Here, each element of the ragged array is assigned a group that then stores the corresponding data array(s) (Fig. 4a). For large ragged arrays this approach is costly as the number of groups and datasets required grows linearly with the number of elements. This approach also hinders efficient search and collective and parallel I/O operations across array components. As performance is a concern for many NWB:N user teams, this motivated the creation of the following alternative design.

#### Region-based Ragged Arrays

In this approach we first concatenate the elements of the ragged array—each of which is a variable-length vector or n-d-array with variable-length first dimension—and store them as a single dataset (Fig. 4b, `<VectorData>`). A second, one-dimensional integer index array then stores for each element of the ragged array the stop-index in the value array, defining the region in the value array that contains the relevant data subsets (Fig. 4b, `<Vectorlndeχ>`). Finally, we define the attribute `target` on the index array, which stores an object reference to the value array to explicitly describe the relationship between the index and value array. This approach supports storage of ragged arrays of arbitrary size using only two datasets and enables efficient collective and parallel I/O and more efficient data search. An alternative approach to using stop indices would be to store region-references as part of the index vector. We opted for stop indices in NWB:N 2.0 because they are more compact and to ease introspection of the indices.

**Figure 5.**
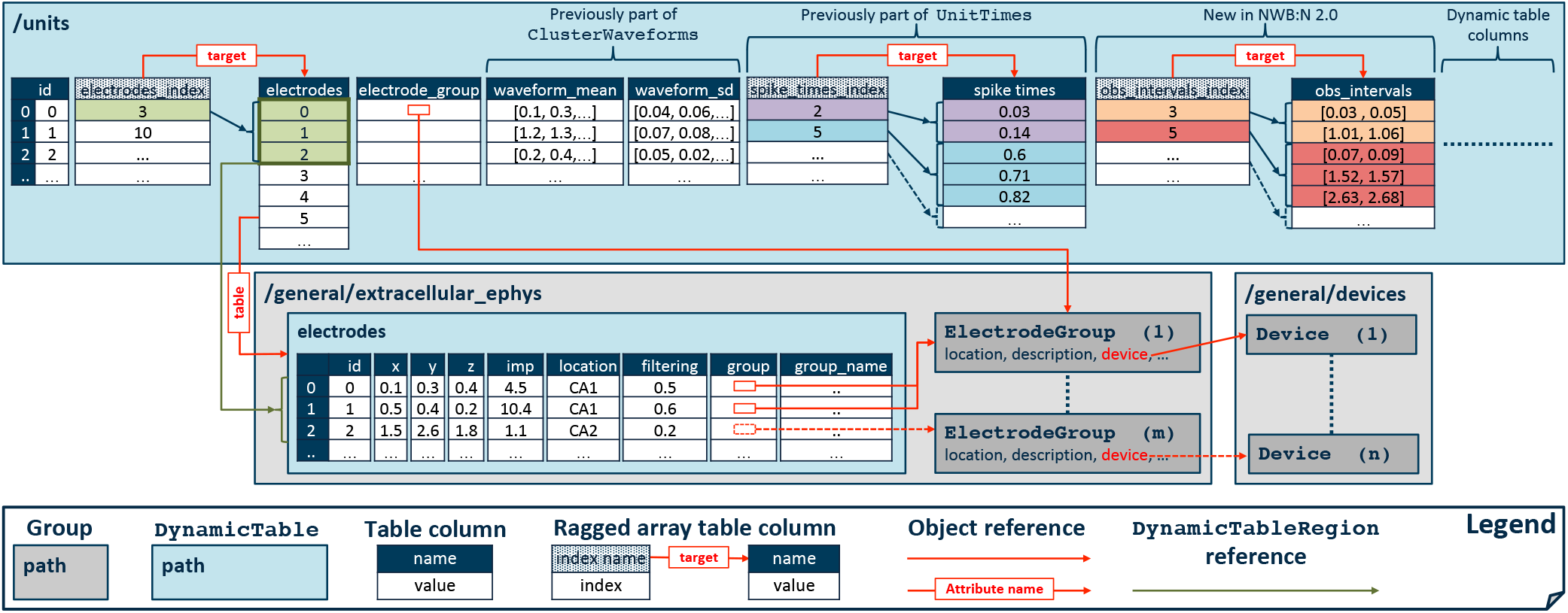
Storing spiking unit data and metadata in NWB:N 2.0. Using a single column-based table allows us to consolidate and greatly simplify the data organization. Using the concept of ragged arrays then supports storage of variable-length spike times and observation intervals directly as part of the units table. Combining the concepts of ragged arrays and row-based table references, then enables referencing of a variable number of electrodes in the electrodes table for each unit.

### 3.6 Integrating Data Tables and Ragged Arrays

A column-based `DynamicTable` is a collection of `VectorData` datasets, each representing a table column. We can, hence, interpret the `VectorData` dataset of a ragged array as a table column that is segmented by the `Vectorlndex` (Fig. 4b). Using this powerful approach enables us to store variable-size data arrays as elements of table columns. As just one example, this approach is useful in the context of feature extraction (e.g., the detection of spiking units in electrophysiology or ROIs in optophysiology) to describe in a single table both feature metadata (e.g., id or location) and associated feature vectors (e.g., the times of neuron spikes or ROI transients). Note, while this approach requires two datasets in HDF5, APIs (e.g., PyNWB) will typically abstract this behavior and instead describe the data as a single table column of variable-length arrays.

## 4 Results

Here we describe the application of the above methods, as well as a broad range of other advances in the NWB:N 2.0 schema and language, to improve storage and management of common neurophysiology data and metadata. In Sec. 4.1 we discuss the application of data tables, ragged arrays and data references to improve organization of neurophysiology data and metadata in NWB 2.0. Afterwards we discuss how refinement to base types has helped to improve organization of acquisition and processing data in NWB:N 2.0 in Sec. 4.2. Sec. 4.3 then presents advances in the schema aimed at improving specificity of the schema and identifiability of objects and types. Finally, in Sec. 4.4 we discuss changes aimed at improving separation of concerns in the NWB:N ecosystem related to the standard schema.

### 4.1 Improving Data Organization via Tables, Ragged Arrays, and Explicit Data References

The combination of tables, ragged arrays, and explicit data referencing allowed us to significantly improve organization and management of electrodes (Sec. 4.1.1), spiking units (Sec. 4.1.2), ROIs (Sec. 4.1.3), sweeps (Sec. 4.1.4), time interval data and metadata (Sec. 4.1.5), and spectral decomposition results (Sec. 4.1.6).

#### 4.1.1 Electrode Metadata

As shown in Fig. 5 (gray boxes), in NWB:N 2.0 we have consolidated all data about individual electrodes in a dynamic, column-based `electrodes` table. To ease data access and management and avoid data duplication, metadata about groups of electrodes is then stored using the extensible type `ElectrodeGroup` and referenced from the electrodes table via explicit object references. Compared to NWB:N 1.0.x, this structure supports: **1. explicit data references** avoiding implicit, text-based references, **2. per-electrode metadata** for impedance and filtering settings, **3. dynamic electrode metadata** via support for dynamic table columns, and **4. efficient data access, search, and maintenance** through the use of a consolidated, standard data table. The chosen structure further avoids the need for per-electrode groups and datasets, ensuring that the number of data objects required remains constant as the density of electrode-based neural recording devices continues to grow.

#### 4.1.2 Spiking Units in Electrophysiology

In NWB:N 1.0.x data about extracted spiking units (i.e., putative single neurons) was stored across several types, specifically: i) `UnitTimes` to store event times of observed units and ii) `ClusterWaveforms` to store waveform shape statistics of clusters. This organization had several critical shortcomings with regard to efficiency, adaptability, and usability.

##### Efficiency

As illustrated in Fig. 4a, previously each unit was stored as a separate group `unit_n` with the times and `description` for the corresponding unit. For use cases with very large numbers of units (e.g., neural simulations or very high-channel-count electrophysiology), this structure required the creation of a very large number of data objects and small, independent I/O operations, reducing I/O and data processing performance and usability. To address this challenge, NWB:N 2.0 organizes the variable-length spike time vectors associated with each unit using the region-based ragged array design described in Sec. 3.5. This approach enables storage of arbitrary numbers of units using a constant number of data objects, supports collective I/O, and eases usability by enabling users to access all units via iteration over a single dataset.

##### Adaptability

A common need is the ability to provide experiment-specific metadata associated with each unit. To avoid the need for custom schema extensions for every experiment, NWB:N 2.0 introduces `/units`, a column-based, dynamic table that enables users to store metadata about units in a convenient and easy-to-use data table.

##### Usability

Finally, to improve usability, we removed the `UnitTimes` and `ClusterWaveforms` types and integrated the corresponding data with the units table. In this way, NWB:N 2.0 makes all unit-specific data accessible via a single, consolidated data table. Fig. 5 illustrates the new structure for storing unit data in NWB:N 2.0.

#### 4.1.3 ROIs in Optophysiology

Image segmentation is used in optophysiology techniques, such as calcium imaging, to extract regions-of-interest (ROIs) describing discrete units of investigation (e.g., neurons). Subsequent extraction of fluorescence traces for each ROI then results in the creation of timeseries describing the activity of each ROI-based unit.

As part of our work with early adopters and through community feedback, we identified several critical issues in NWB:N 1.0. x. **1. Efficiency:** Similar to the organization of spiking units, efficiency and usability were a concern as NWB:N 1.0. x required the creation of separate groups and datasets for each individual ROI. **2. Implicit Relationships:** Relationships between ROIs and the corresponding time series (i.e., `ROIResponseSeries`) were only defined implicitly via a string, so that users had to know *a priori* which `ImageSegmentation` and `ImagingPlane` was used to produce the ROI. **3. 3D ROIs:** Finally, lack of support for three-dimensional ROIs produced by multi-plane imaging was identified as a central issue for users.

**Figure 6.**
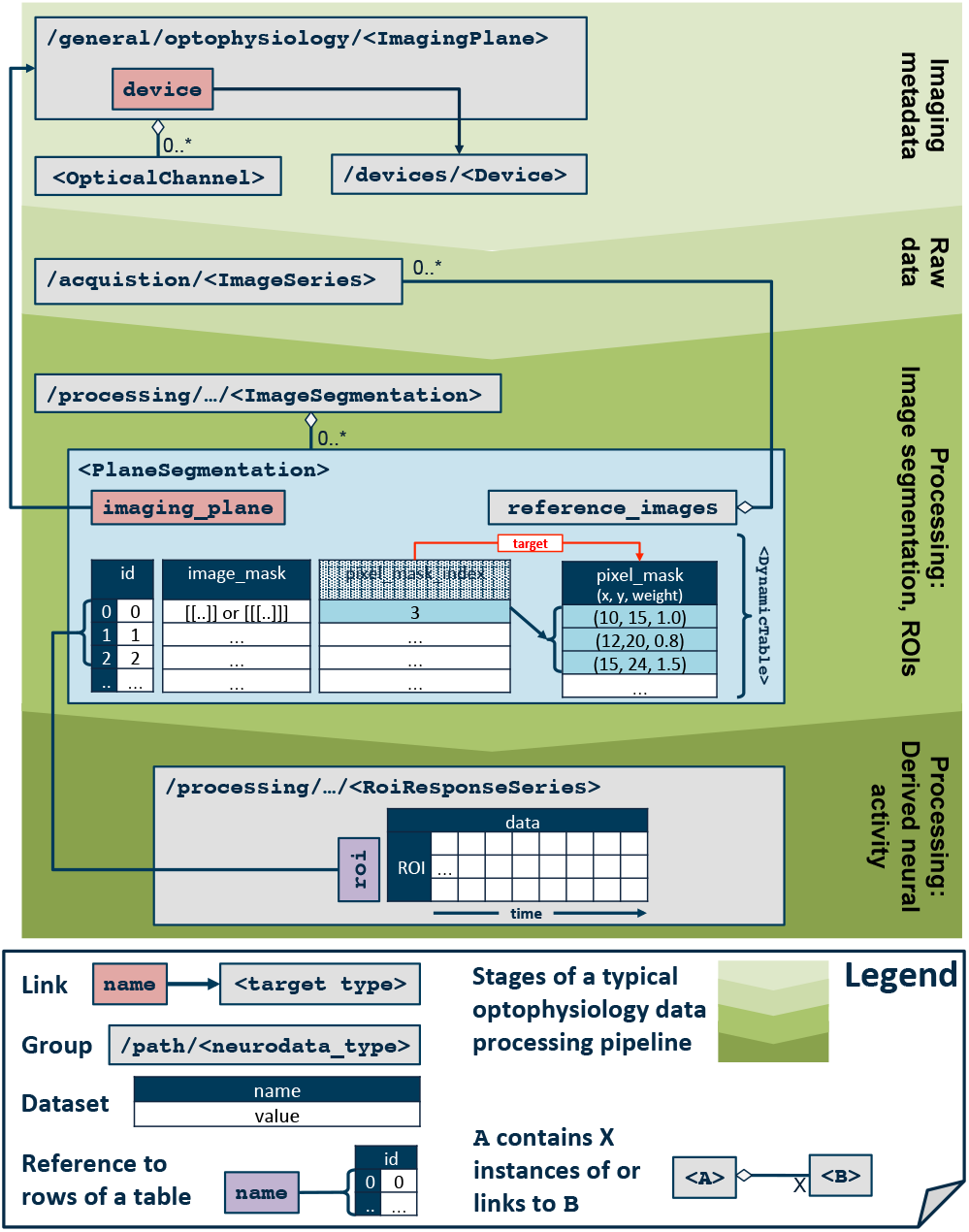
Overview of the main data types used in the stages of a typical optophysiology analysis pipeline.

To address these needs, NWB:N 2.0 uses dynamic, column-based tables to efficiently store large collections of ROIs in combination with explicit data references to ensure unique identification of related objects (Fig. 6). To support 2D and 3D ROIs the `PlaneSegmentation` table supports both pixel and voxel masks as well as 2D and 3D image masks. For a detailed list of changes compared to NWB:N 1.0.x see the online release notes^11^.

#### 4.1.4 Sweeps

In intracellular electrophysiology it is common to have sweeps, similar to trials. This results in the need to associate multiple time series with each other, or more specifically, to group a set of `PatchClampSeries`. NWB:N 1.0.x did not support the concept of sweeps. To address this need, NWB:N 2.0 adds a dynamic, column-based `SweepTable` to the `/general/intracellular_ephys` group used to store intracellular electrophysiology data. The sweep table stores for each sweep a unique sweep number as well as a variable-length vector of patch-clamp time series associated with the sweep. The latter uses the concept of ragged arrays integrated with data tables described earlier in Sec. 3.6.

**Figure 7.**
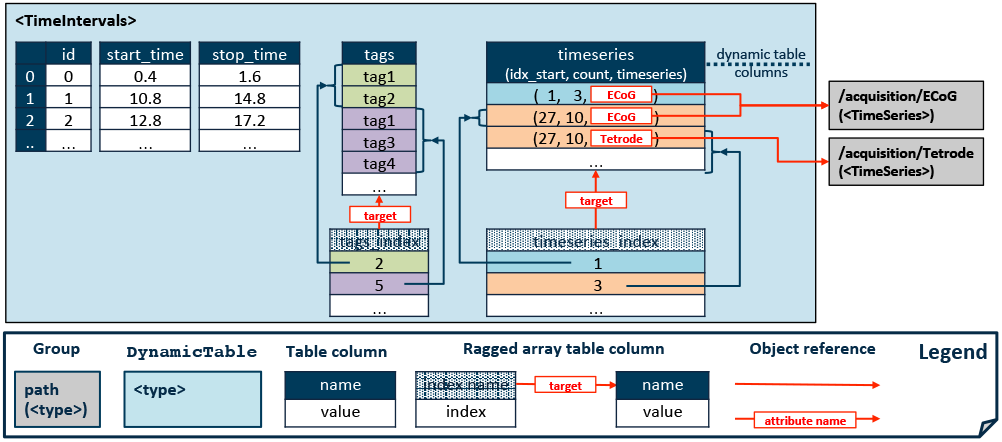
Storing time interval tables in NWB:N 2.0. See Fig. 2 for an example specification of the compound dataset “timeseries” using the NWB:N specification language.

#### 4.1.5 Managing Time Intervals

A common need in neurophysiology is the ability to describe time intervals, i.e., ranges in time associated with particular events or features, e.g., spikes, experimental intervals, stimuli or behavioral events, among many others. To address this need, NWB:N 1.0.x specifically supported the concept of epochs, where each epoch was stored as a separate group as part of the top-level `/ epochs` group.

While this concept was generally useful, we identified three critical unmet needs as part of our requirements analysis and from user feedback. **1. Efficiency:** Users often need to store very large numbers of epochs (and other time intervals). This led to concerns with regard to efficiency and usability due to the need for individual groups/dataset for each epoch in NWB:N 1.0.x. **2. Adaptability:** Users commonly need to store additional metadata for time intervals. With the removal of “Custom” to allow users to arbitrarily add data, this means that using the original design required users to create format extensions to add custom metadata. **3. Generality:** Beyond experimental epochs, users often require collections of different kinds of time interval data, each with different sets of metadata. Specifically, lack of support for trials was identified as a common issue for users.

To address these needs, we have generalized the concept of storing collections of time intervals. Specifically, NWB:N 2.0 defines the new type `Timelntervals`, a dynamic, column-based table for storing time intervals (Fig. 7). For each time interval (i.e., row in the table) we store an id, start time, stop time, and (using the concept of region-based ragged array table columns) a variable length list of tags and timeseries references. More specifically, the “timeseries” column is a compound dataset that stores the object reference to the `TimeSeries` object along with the corresponding start index (`idχ_start`) and number of timesteps (`count`) that the specific start and stop times map to. This approach allows us to define general, dataset-independent time intervals while at the same time supporting direct look-up of data in related time series. By storing time intervals in a dynamic table, the `Timelntervals` type addresses (1) the need for more efficient and convenient storage by allowing users to store large collections of time intervals in a convenient table as well as (2) the need for adaptability by supporting the dynamic addition of table columns.

To support (3) the need for experiment trials and the ability to store multiple collection of time intervals, we introduced a new top-level group `/intervals`. Here users can store arbitrary collections of `Timelntervals` tables, including the predefined tables for epochs, trials, and invalid times.

#### 4.1.6 Spectral Decomposition

A common analysis in neurophysiology is to decompose measured neural or behavioral signals using frequency decomposition or bandpass filtering. NWB:N 2.0 introduces a new time series type `DecompositionSeries` to offer a common interface for labs to exchange the result of common time-frequency analysis. The series includes a link to the source `TimeSeries` object to support unique identification of the raw signal that is being decomposed. The metadata needed to fully describe the bands often depend on lab needs and the decomposition method used. To address this need, we use a dynamic, column-based table with standard columns for band name, limits, and (optionally) mean, and standard deviation while allowing users to flexibly add new band features without the need for custom format extensions.

### 4.2 Refining Base Types to Improve Storage of Acquisition- and Processing Data

NWB:N 2.0 makes the storage of acquisition data (Sec. 4.2.1) and results from standardized processing pipelines (Sec. 4.2.2) more flexible and easy to use.

#### 4.2.1 Acquisition Data

NWB:N 1.0.x defined the group `/acquisition` for storage of acquisition data, which included subgroups for `timeseries` and `images`. Here we identified two main challenges. First, the limitation to only allow timeseries as acquisitions was too restrictive for many users. Second, the fixed structure of the subgroup for images was not reusable and specific enough to support the common needs in neurophysiology for storing additional photographs and other imagery related to an experiment.

To improve flexibility, NWB:N 2.0 allows users to store any primary data type (i.e., `NWBDatalnterface`, Sec. 3.1) as part of the acquisition group and removed the timeseries and images subgroups. Also, to improve specificity and reusability for storage of additional images, NWB:N 2.0 adds the generic dataset type Image and corresponding subtypes for `Grayscalelmage`, `RGBImage`, and `RGBAImage` as well as a corresponding `NWBDatalnterface` container type `Images` for storing basic image collections. Note, in addition to these generic image types for storing arbitrary image collections, NWB:N provides dedicated types for storing timeseries of images from optophysiology experiments.

#### 4.2.2 Processed Data

Results from standardized processing pipelines are stored in `ProcessingModules` as part of the top-level `processing/` group in NWB:N.

##### ProcessingModule

NWB:N 1.0.x defined the concept of a “Module,” which is a container for storing results from common data processing pipelines. In NWB:N 2.0, “Module” has been renamed to the more specific term `ProcessingModule` to avoid ambiguity and help clarify the purpose and use of this type. Also, similar to the acquisition group, a processing module may now contain any `NWBDataInterface`, which, as we will discuss next, has allowed us to significantly improve flexibility in the definition of processing modules.

##### NWBDataInterface

NWB:N 1.0.x introduced the type “Interface” which was used as the base type that all objects in a processing module had to inherit from. This design led to several challenges in practice. First, data products from preprocessing and analyses are often derived time series. However, as “Interface” was not a base type of `TimeSeries`, this meant that analysis containers could not directly inherit from `TimeSeries` (but could only contain them) and that processing modules could not directly contain timeseries objects. Second, all analysis types in NWB:N 1.0.x had fixed names so that each processing module could only contain a single instance of each analysis. Third, the name “Interface” was seen as ambiguous and often led to confusion among users and developers as towards the function and purpose of the type.

To address these challenges, `NWBDataInterface` was introduced as a more general base type (Sec. 3.1) and replacement for “Interface”, and now serves as a base type for all primary data (including `TimeSeries`). This generalization, while seemingly simple, has been critical to enable users to create analysis data containers that extend `TimeSeries` and to permit storage of timeseries types directly as part of processing modules. Further, NWB:N 2.0 now defines default names (rather than fixed names) for all analysis types so that users can store multiple instances of the same analysis in a processing module and customize their names. Finally, the renaming and generalization of “Interface” has helped to clarify the purpose and use of the new `NWBDataInterface` type.

### 4.3 Improving Specificity and Identifiability

To enable more accurate data interpretation, NWB:N 2.0 improves the specificity of reference timestamps (Sec. 4.3.1), ensures that all objects can uniquely be identified (Sec. 4.3.2), makes links explicit (Sec. 4.3.3), supports specification of data shapes (Sec. 4.3.4) and default names and values (Sec. 4.3.5), and includes various other enhancements to improve consistency and use (Sec 4.3.6).

#### 4.3.1 Defining Reference Time

NWB:N 1.0.x defined the start time of the session as the global clock that all timestamps in a file are synchronized with. However, the format for storing the reference time was not defined, allowing users to store the session start time as an arbitrarily formatted string. To address this need, we extended the specification language and APIs to support ISO8061 formatted datetimes^2^ as the datatype `isodatetime`, which allows us to enforce the use of ISO8061 in the NWB:N 2.0 schema, ensuring consistent human and programmatic interpretation of timestamps within and across data files.

In addition, the use of session start time to indicate both 1) the start time of a session and 2) the reference clock for all timestamps in a file meant that users could only store times relative to a session. Relative times are commonly used to support direct interpretation and analysis of individual sessions. However, use cases involving files from many sessions or subjects, and more broadly management of data across neuroscience labs and projects, benefit from having the ability to define a common standard reference time across files (e.g., `1970-01-01T00`: `00`: `00Z` in the case of POSIX time). To support these use cases, NWB:N 2.0 now defines a separate timestamp reference time in addition to the session start time. For convenience, the reference time is by default set to the same value as the session start time (assuming common relative times).

#### 4.3.2 Ensure Unique Object Identifiability

NWB:N 1.0.x allowed untyped groups and datasets with user-defined names as part of the schema. In the case where multiple objects with variable name and no assigned type are contained in the same group, this leads to ambiguity with regard to which part of the schema governs a particular object. To ensure that each object can be uniquely associated with the corresponding description in the schema, NWB:N 2.0 requires that each dataset and group must either have a unique type (i.e., neurodata_type) or fixed name.

To ensure compliance with this rule, we defined previously missing neurodata_types for a number of types, e.g., `ImagingPlane`, `IntracellularElectrode`, or `OptogeneticStimulusSite` among others. By defining a unique type for previously untyped schema also helped improve reuse and extensibility of types, e.g., to facilitate extension for `Subject` metadata. Finally, to ensure that we can uniquely identify the schema for each object in an HDF5 file, we defined a set of reserved attributes, which are populated automatically based on the schema. These attributes store the neurodata_type and information about the name and version of the namespace where the type is defined.

#### 4.3.3 Making Data Links Explicit

The original NWB:N 1.0.x schema specified a number of datasets containing implicit links, i.e., datasets with lists of either 1) strings with object names, 2) strings with paths, or 3) integer indexes to implicitly point to other locations in an NWB:N file. These forms of implicit links were not self-describing, e.g., the kind of linking, target location, implicit size and numbering assumptions could not easily be identified. As such, this approach hindered human interpretation of the data as well as programmatic resolution of these kind of links. In the NWB:N 2.0 schema we have modelled these implicit relationships explicitly through a combination of reorganization of metadata using tables and the use of links and datasets of object- and region-references. See Sec. 4.1 and Appendix D for an overview of corresponding changes in the schema.

#### 4.3.4 Specifying Data Shape

In addition to labels for dataset dimensions via the dims key (a.k.a. “dimensions” in NWB:N 1.0.x), the NWB:N 2.0 specification language adds the key `shape` to support restriction and validation of the shape of datasets. Specifying the shape of datasets has helped to clarify and further formalize the specification of a large number of datasets in the NWB:N 2.0 schema.

#### 4.3.5 Supporting Default Names and Values

In addition to specification of fixed names and values, NWB:N 2.0 adds the ability to define default names for groups and datasets and default values for datasets and attributes. This capability is useful to simplify use of the NWB:N data standard by providing users with practical defaults wherever possible while still enabling users to customize the values if necessary.

#### 4.3.6 Others: Consistency and Identifiability

Beyond the above main schema changes, NWB:N 2.0 implements a broad range of additional schema changes to improve consistency and identifiability. For example, names have been harmonized and missing metadata fields added in the schema to improve consistency of object specifications across types. Several previously required groups (e.g., for epochs and optogenetics metadata) have been made optional to reduce the need for empty groups. Also, NWB:N 2.0 adds a keywords field to allow users to define keywords for their data to ease integration with data archives. NWB:N 2.0 further adds the type `LabMetaData` as part of `/general`. This allows users to more easily integrate lab-specific metadata by extending the LabMetaData type.

### 4.4 Improving Separation of Concerns

One main goal during the development of NWB:N 2.0 was to identify, define, and separate the main components of the NWB:N ecosystem. This process ultimately also required changes to both the data standard schema (Sec. 4.4.1) as well as the specification language (Sec. 4.4.2) to improve management and improve separation of concerns.

#### 4.4.1 Refine and Remove Common Attributes

Here we discuss changes to common attributes in the NWB:N schema that have been removed or refined in NWB:N 2.0.

##### ancestry

In NWB:N 1.0.x the attribute `ancestry` was used to explicitly store the list of types a given type inherits from with the goal to allow users to easily inspect the ancestry of an object. As the type hierarchy is encoded in the schema, the ancestry attribute stored redundant information, which lead to maintenance issues as well as possible inconsistencies in the schema, as the ancestry attributes of every object had to be updated any time the type hierarchy changed in the schema. As such, the ancestry attribute has been removed in NWB:N 2.0. Inspection of the ancestry of a type or object is ultimately the function of the data and specification APIs and much more appropriately solved there.

##### neurodata_type

Similar to ancestry, NWB:N 1.0.x explicitly defined the attribute “neurodata_type” to store the type of an object. However, as the type of an object is defined by the neurodata_type key in the schema (not the attribute object), this approach easily lead to inconsistencies between the stored and actual type. As such, the manually defined attribute neurodata_type has been removed for all types in the NWB:N 2.0 schema. Instead NWB:N 2.0 defines a corresponding reserved attribute that is automatically populated with the value of the “neurodata_type” key in the schema.

##### Custom

NWB:N 1.0.x permitted users to add arbitrary data to the file simply by defining the neurodata_type attribute of a group or dataset in a file as Custom. Unfortunately, this approach makes it hard to interpret the data, design effective APIs, and interact with the data in a standard fashion. One main decision in the design of NWB:N 2.0 has been that all objects stored in the file must be governed by the schema. As such, NWB:N 2.0 requires the use of schema extension for integration of new data types and support for dynamic addition of objects with “neurodata_type=Custom” has been removed. For metadata, NWB:N 2.0 provides the added flexibility via support for dynamic columns in tables. This approach ensures that all objects in a file are indeed governed by the schema while at the same time providing well-defined mechanisms for dynamic integration of experiment-specific metadata.

##### source

In NWB:N 1.0.x the attribute `source` was defined as a free text attribute intended for storage of provenance information. Practical experience with early adopters, however, consistently showed that instead of encoding provenance, the source attribute was often either ignored, contained no useful information, or was misused to encode custom metadata. The source attribute has, therefore, been removed as a required field from the base types in NWB:N 2.0 (Sec. 3.1). However, the NWB:N community recognizes data provenance as a critical issue and we plan to investigate more advanced options for integration of data provenance with NWB:N as part of our future work.

#### 4.4.2 Specification Language

As part of the development of NWB:N 2.0 we have overhauled the specification language significantly. Here we focus on two main changes that affected the data standard schema.

##### autogen

In NWB:N 1.0.x the key `autogen` was used to specify computations in the schema that an API would then have to carry out to automatically create datasets and attributes derived from other fields. This feature was implemented in response to requirements voiced during the early development of NWB:N 1.0.x that data files should be human-readable using the HDFView utility. With the emergence of PyNWB and MatNWB as dedicated NWB:N data APIs, this requirement has since been dropped. After a careful review of `autogen` at the 3rd NWB:N hackathon at HHMI it was decided to remove the `autogen` feature and all corresponding objects from the NWB:N 2.0 schema. See Appendix C for details.

##### Separating keys and values

One main focus of our work on the specification language as part of the NWB:N 2.0 efforts was to ensure that all information is being defined explicitly via dedicated key/value pairs. This requirement helped to significantly improve readability of the schema, specificity and interpretability of the schema, and avoids potential collisions between keys. For example, in NWB:N 1.0.x the specification of each object was identified in the schema language by a key constructed from a regular expression of a combination of the path, name, type and quantity of the corresponding specification. This led to keys that were hard to read and construct, implicitly encoded specification values in keys, and ultimately led to collision of keys. In NWB:N 2.0 the schema language has been updated to define explicit key/value pairs for all specification details, e.g., groups, datasets, links, name, dtype, or quantity among others. For details please see the specification language documentation^7^.

##### Simplify Reuse of Types

The concept of a neurodata_type is similar to the concept of a class in object-oriented programming. A neurodata_type is a unique identifier for a specific type of group or dataset in the schema. Assigning a neuro-data_type enables the reuse of the specified type by inclusion or inheritance. Previously, in NWB:N 1.0.x reuse of types was implemented via a combination of multiple different keys, specifically, `merge`, `merge+`, `include`, `properties`, and `allow_subclasses`^12^.

To simplify the specification and reuse of types, the NWB:N 2.0 specification language instead defines the keys: **1) `neurodata_type_def`** to define (i.e, create) a new type and **2) `neurodata_type_inc`** to define the base type of a type. The combination of neurodata_type_inc and neuro-data_type_def provides an easy-to-use mechanism to reuse types via both inheritance (i.e., extension of a type) and composition (i.e, embedding of a type as a component of a new type) as follows:

1. **Create type:** If only neurodata_type_def is set then we create a new type from scratch without a base type.
2. **Include type:** If only neurodata_type_inc is set then we include (reuse) an existing type.
3. **Extend type:** If both neurodata_type_inc and neuro-data_type_def are set, then we create a new type that inherits from an existing type.
4. **Create named object:** Finally, if neither neuro-data_type_inc nor neurodata_type_def are defined then we define a standard dataset or group without a type and, hence, the object must have a fixed name to ensure the object can be uniquely identified.

## 5 Conclusions

### Community

A central goal of NWB:N 2.0 is to work with the neuroscience community towards better science solutions. User, developer and application engagement are central to this mission. In April 2018, 65+ scientists from 20 major institutions attended the NWB:N development and user hackathons at AIBS and LBNL. Also, many groups are already adopting NWB:N, including laboratories at UCSF, UCB, Caltech, LBNL, HHMI Janelia Farm, and the Allen Institute for Brain Science, among others. The NWB:N teams are also actively engaging with NIH BRAIN Initiative projects, and the Kavli foundation funded several seed grants to help various labs to evaluate and adopt NWB:N. Also, engagement with industry partners, e.g., Kitware (visualization, continuous integration), Vidrio (MatNWB), MathWorks (Matlab), Vathes (data management) and others, is central to our software strategy and to facilitate the creation of an advanced data science ecosystem for neurophysiology.

### Governance

As an open data standard, all software, documentation, communication channels, and other NWB:N resources are open to the community via common tools, e.g., GitHub, Slack, GoogleGroups, ReadTheDocs, PIP or Conda, and accessible via https://neurodatawithoutborders.github.io. Development on NWB:N is currently funded via several projects by the Kavli foundation and NIH. The NWB:N Executive and Technical Advisory Boards together with our sponsors and project teams, play a central role in creating the vision for and enabling NWB:N to grow to become a widely accepted standard in neurophysiology.

### Future Directions

With the release of NWB:N 2.0, the next main phase of the NWB:N project begins. One main goal is to promote adoption of NWB:N 2.0 and to transition the standard into production use in neuroscience laboratories, projects, and data archives. Another main goal is to develop NWB:N 2.x to enhance the standard for new applications and use cases.

To facilitate integration with lab data management, we plan to develop methods and tools to support “foreign fields” to enable referencing of data stored in other systems directly from NWB:N files. This approach promises to enable users to seamlessly link data across systems, easily access linked data via familiar NWB:N file mechanisms, avoid duplicate storage of data across systems, and integrate NWB:N with lab data management.

To enable efficient discovery of data, it is critical that metadata is specified using interpretable and consistent terms. To address this challenge we will integrate standardized metadata models, controlled vocabularies, and ontologies with NWB:N.

An often neglected but critical need in data-intensive analyses is the ability to efficiently identify, select, and process relevant data subsets. To address this challenge, we will develop tools to make it easy to query and analyze NWB:N data in parallel using modern commodity and high-performance compute systems. More broadly, the goal is to develop easy-to-use data analytics and visualization tools and libraries for NWB:N as well as integration of NWB:N with existing neurophysiology codes.

Finally, to enable the neuroscience community to adopt and curate the NWB:N standard and facilitate the integration of new use-cases, we will develop standard mechanisms for extension and tools and processes for sharing, evaluation, acceptance, and use of NWB:N extensions.

## Acknowledgements

This work was sponsored by the Kavli foundation. Research reported in this publication was supported by the National Institute Of Mental Health of the National Institutes of Health under Award Number R24MH116922 to O. Ruebel, and Award Number 5R44MH115731-02 to W. Schroeder, and supported by the BRAIN Initiative under award number U19-NS104590 to I. Soltesz, and by the Simons Foundation for the Global Brain grant 521921 to L. Frank, and by additional funding from the Simons Foundation for MatNWB. The content is solely the responsibility of the authors and does not necessarily represent the official views of the National Institutes of Health.

We thank the participants of the 3rd, 4th, and 5th NWB:N hackathons and the NWB:N user and developer community for their feedback and contributions. We thank the members of the NWB:N Executive Committee Katrin Amunts, Kris Bouchard, Loren Frank, Christof Koch, Markus Meister, Friedrich Sommer and Karel Svoboda.

## Author contributions statement

**O. Rübel** is the lead for the NIH Award R24MH116922 for development of NBW:N, documentation lead, and contributed to both the NWB:N 2.0 APIs and schema. **A. Tritt** is the lead software architect of PyNWB and led many of the schema changes described in the manuscript. **B. Dichter** contributed to MatNWB, PyNWB, the decomposition series and many other schema changes and developed NWB:N integration and conversion codes and strategies with neuroscience groups. **T. Braun** contributed to PyNWB as well as the schema as part of IS08061 adoption, SweepTable, and refinement/removal of common attributes among others. **N. Cain** contributed to PyNWB, the NWB:N schema, use case testing, and user training. **N. Clack** contributed as a core developer to MatNWB. **T.J. Davidson** contributed to requirements, use cases, and evaluation of the schema and API. **M. Dougherty** contributed to requirements, use cases, and evaluation of PyNWB. **J.-C. Fillion-Robin** contributed to the design and implementation of continuous integration and open source software practices. **N. Graddis** contributed to PyNWB, the NWB:N schema, and use-case testing. **M. Grauer** contributed to the implementation of continuous integration and open source software practices. **J. Kiggins** contributed to use cases, requirements, and evaluation of PyNWB and schema features. **L. Niu** contributed as a core developer to MatNWB. **D. Ozturk** contributed to the implementation of continuous integration and open source software practices. **W. Schroeder** is the PI for NIH Award 5R44MH115731-02 and contributed to project and open source community leadership. **I. Soltesz** was the lead PI of the U19-NS104590 award, and provided input, feedback and financial support for the project. **F. Sommer** lead the first NWB:N pilot project and provided input and feedback during the development of NWB:N 2.0. **K. Svoboda** provided input and feedback during the development process and provided financial support for MatNWB. **L. Ng** is co-lead for the NIH Award R24MH116922 and SBIR with Kitware and contributed to project leadership and use cases. **L. Frank** provided input and feedback during the development process and provided financial support for the project. **K. Bouchard** was lead PI of the NWB4HPC project and provided input and feedback during the development process. All authors reviewed the manuscript.

## Disclaimer

This document was prepared as an account of work sponsored by the United States Government. While this document is believed to contain correct information, neither the United States Government nor any agency thereof, nor the Regents of the University of California, nor any of their employees, makes any warranty, express or implied, or assumes any legal responsibility for the accuracy, completeness, or usefulness of any information, apparatus, product, or process disclosed, or represents that its use would not infringe privately owned rights. Reference herein to any specific commercial product, process, or service by its trade name, trademark, manufacturer, or otherwise, does not necessarily constitute or imply its endorsement, recommendation, or favoring by the United States Government or any agency thereof, or the Regents of the University of California. The views and opinions of authors expressed herein do not necessarily state or reflect those of the United States Government or any agency thereof or the Regents of the University of California

## Appendix

### A Specifying Data Tables

Here we provide abbreviated example specifications for the tables shown in Fig. 3 using the NWB:N specification language. Fig. A.1 shows a row-based table in which each row is stored as a single element of a 1D compound-type array. Here, the columns of the table are described by the components of the compound data type. In contrast, the column-based table shown in Fig. A.2 stores each column as a separate dataset, i. e., each row spans multiple datasets. Fig. A.3 then illustrates a hybrid approach in which the (*x, y, z*) location is stored in a single compound data-type column as part of an otherwise column-oriented table.

**Figure A.1.**
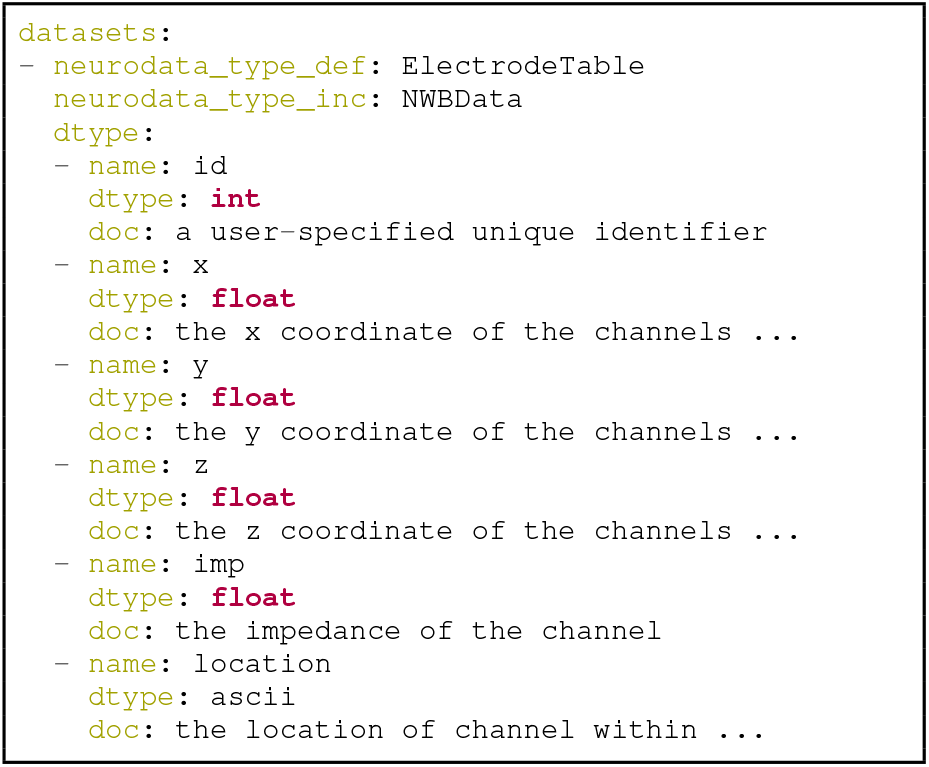
Abbreviated specification of a row-based table specified via a 1D compound-type array, for storing electrode metadata (see Fig. 3a).

**Figure A.2.**
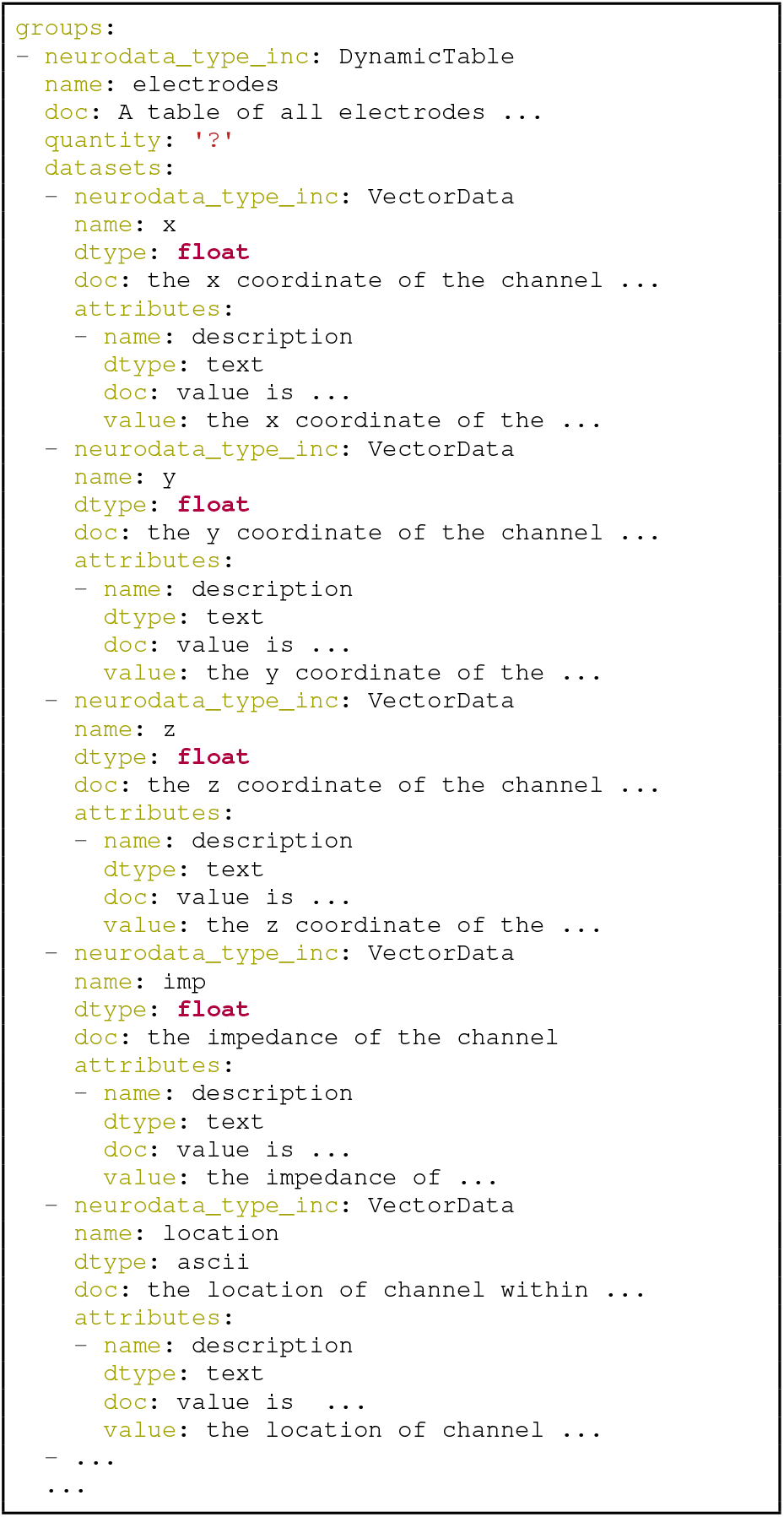
Abbreviated specification of a column-based table for storing electrode metadata (see Fig. 3b). Note, the id column is inherited from `DynamicTable`.

**Figure A.1.**
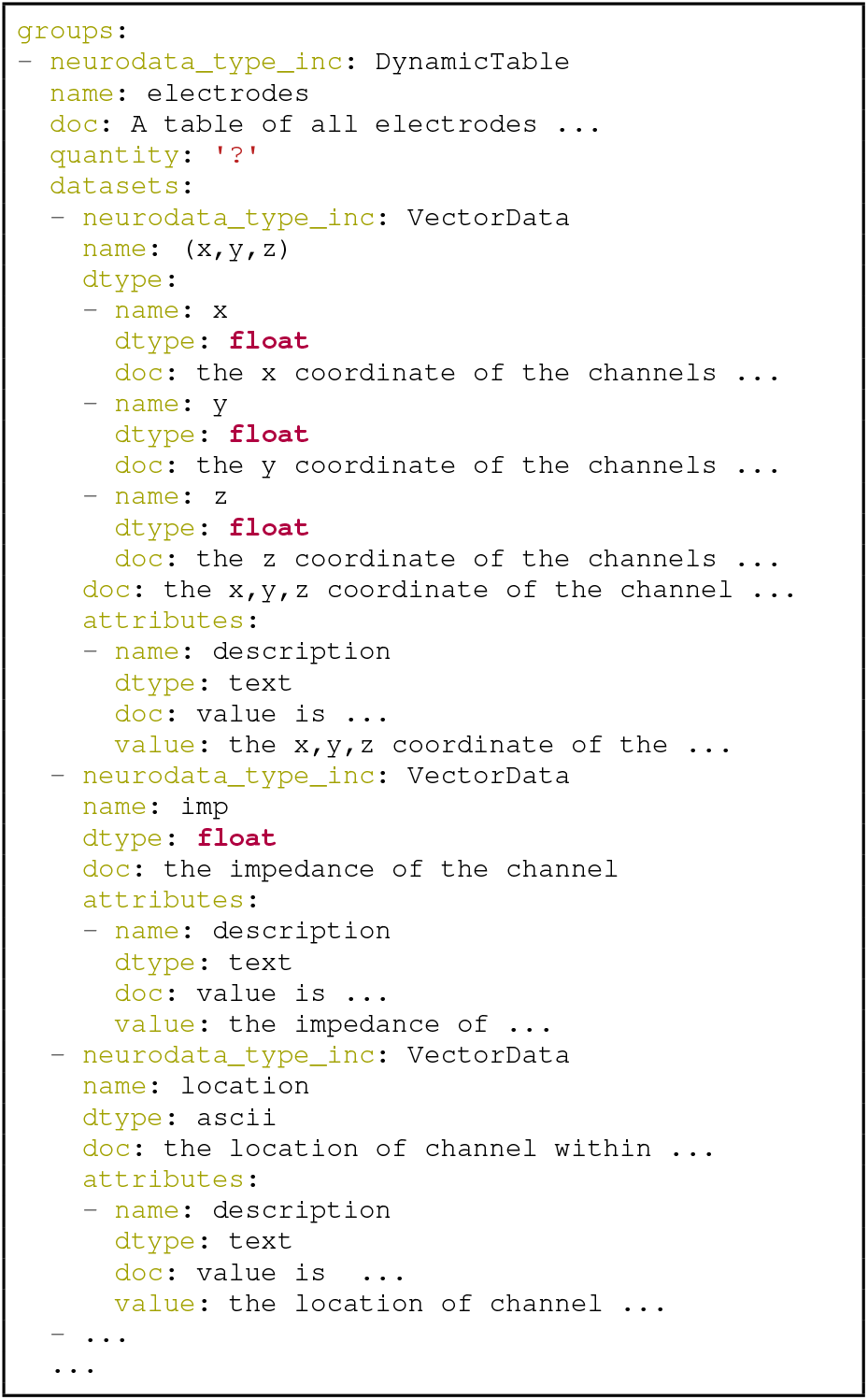
Abbreviated specification of a hybrid table for storing electrode metadata (see Fig. 3c). Note, the id column is inherited from `DynamicTable`.

### B Dense Ragged Arrays

As an alternative to the approach of implicit ragged arrays and region-based ragged arrays described in Sec. 3.5, NWB:N generally also supports dense ragged arrays. Using a single dense array, we can represent ragged arrays by assigning a unique default value (e.g., *NaN*) to unused entries and as such, regularize the array (Fig. B.1). Depending on the raggedness of the array, this approach can lead to potentially large storage overheads due to the allocation of unused data values. Here, storage optimizations like chunking and compression can help reduce storage overheads but may also introduce additional memory and I/O cost. While NWB:N generally supports this approach to store ragged arrays, the NWB:N 2.0 data standard currently does not use it due to useability and cost concerns.

**Figure B.1.**
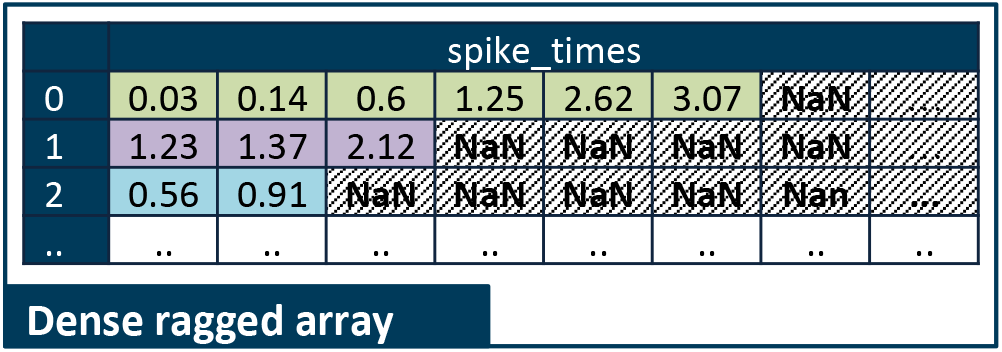
Illustration showing the use of a dense arrays to model a ragged array storing the times of spikes from single units. See Fig. 4 for an example showing the use of implicit and region-based ragged arrays to describe the same data.

### C autogen

Support for autogen has been removed in NWB:N 2.0. The autogen specification was originally used to specify that the contents of attributes or datasets (i.e,. their values) can be derived from the contents of the HDF5 file and, hence, generated and validated automatically. As such, autogen crossed a broad range of different functions, including:

- Specification of the structure of format datasets and attributes.
- Description of data constraints (e.g., the shape of the generated dataset directly depends on the structure of the input data consumed by autogen).
- Specification of the content (i.e., value) of datasets and attributes.
- Description of computations to create derived data.
- Validation of the structure and content of datasets and attributes.

This mixing of functionality in turn led to several concerns:

- autogen exhibited a fairly complex syntax, which made it hard to interpret and use.
- autogen was specifically used to create derived data from information that is already in the NWB:N file. Attributes and datasets generated via autogen, hence: 1) were redundant, 2) required bookkeeping to ensure data consistency, 3) generate dependencies across data and types, 4) had limited utility as the information could be derived through other means, and 6) interpretation of data values may require the provenance of autogen.
- Description of computations as part of a format specification was seen as problematic.
- There was potential for collisions between autogen and the specification of the dataset or attribute itself.

#### C.1 How did the removal of autogen affect the NWB:N schema?

autogen was used in NWB V. 1.0.6 to generate 17 datasets/attributes primarily to separately store: 1) the path of links or 2) lists of names of datasets and groups of a given type/property. The datasets were reviewed at the 3rd NWB:N hackathon at HHMI and determined to be non-essential and as such removed from the data standard schema as well. Below a list of the datasets and attributes that were generated via autogen in NWB:N 1.0.6 that have been removed in NWB:N 2.0.

1. Datasets and attributes that have been removed due to redundant storage of the path of links stored in the same group:

- `<IndexSeries>/indexed_timeseries_path`
- `<RoiResponseSeries>/segmentation_interface_path`
- `<ImageMaskSeries>/masked_imageseries_path`
- `<ClusterWaveforms>/clustering_interface_path`
- `<EventDetection>/source_electricalseries_path`
- `<MotionCorrection>/image_stack_name/original_path`
- `<NWBFile>/epochs/epoch_X/links`
2. Datasets and attributes that have been removed because they stored only a list of groups or datasets (of a given type or property) in the current group.
  - `<Module>/interfaces`
  - `<ImageSegmentation>/image_plane/roi_list`
  - `<UnitTimes>/unit_list`
  - `<TimeSeries>/extern_fields`
  - `<TimeSeries>/data_link`
  - `<TimeSeries>/timestamps_link`
  - `<TimeSeries>/missing_fields`
3. Other datasets/attributes that have been removed to ease use and maintenance because the data stored is redundant and can be easily extracted from the file:
  - `<NWBFile>/epochs/tags`
  - `<TimeSeries>/num_samples`
  - `<Clustering>/cluster_nums`

#### D Replaced Implicit Links/Data-Structures with Explicit Links

NWB:N 1.0.6 specified datasets of implicit links, containing lists of either 1) strings with object names, 2) strings with paths, or 3) integer indexes to implicitly point to other locations in an NWB:N file. These forms of implicit links were not self-describing (e.g., the kind of linking, target location, implicit size and numbering assumptions were not easily identified). This hindered human interpretation of the data as well as programmatic resolution of these kind of links. Below we provide an (incomplete) list of datasets that have been replaced to make implicit links between data explicit:

- The text dataset `image_plane` in the `TwoPhotonSeries` type has been changed to a link to the corresponding `ImagingPlane` (stored in /general/optophysiology)
- The text dataset `image_plane_name` in the ImageSegmentation type has been changed to a link to the corresponding ImagingPlane (stored in `/general/optophysiology`). The dataset was also renamed to `image_plane` for consistency with the TwoPhotonSeries type.
- The text dataset `electrode_name` in the `PatchClampSeries` type has been changed to a link to the corresponding `IntracellularElectrode` (stored in `/general/intracellular_ephys`). The dataset was renamed to electrode for consistency.
- The text dataset site in the `OptogeneticSeries` type has been changed to a link to the corresponding `StimulusSite` (stored in `/general/optogenetics`).
- The integer dataset `electrode_idx` in the `FeatureExtraction` type has been changed to the dataset `electrodes` of type `DynamicTableRegion` which points to a region of the `ElectrodeTable` (stored in `/general/extracellular_ephys/electrodes`).
- The integer array dataset `electrode_idx` in the `ElectricalSeries` type has been changes to the dataset `electrodes` of type `DynamicTableRegion` which points to a region of the ElectrodeTable (stored in `/general/extracellular_ephys/electrodes`).
- The text dataset `<electrode_group_X>/device` in `/general/extracellular_ephys/` has been changed to the link `<ElectrodeGroup>/device`.
- The text dataset `<electrode_group_X>/device` in `/general/intracellular_ephys/` has been changed to the link `<ElectrodeGroup>/device`.
- Also, as discussed in Sec. 4, the `Epochs`, `Unit`, `Electrodes`, `SweepTable`, `TimeIntervals` and other `DynamicTable` types in NWB:N 2.0 also use region and object references to explicitly reference other data.

